# Design and construction of 3D-printed devices to investigate active and passive bacterial dispersal on hydrated surfaces

**DOI:** 10.1101/2022.05.16.492069

**Authors:** Thierry Kuhn, Matteo Buffi, Saskia Bindschedler, Patrick S. Chain, Diego Gonzalez, Claire Stanley, Lukas Y. Wick, Pilar Junier, Xiang-Yi Li Richter

## Abstract

To disperse in water-unsaturated environments, such as the soil, bacteria rely on the availability and structure of water films forming on biotic and abiotic surfaces, and, especially, along fungal mycelia. Dispersal along such “fungal highways” is driven both by mycelial physical properties and by interactions between bacteria and fungi. To understand the role of abiotic elements, we designed and 3D-printed two devices establishing stable liquid films that support bacteria dispersal in the absence of biotic interactions. The thickness of the liquid film determined the presence of hydraulic flow capable of carrying non-motile cells. In the absence of flow, only motile cells can disperse in the presence of an energy source. Non-motile cells could not disperse autonomously without flow, but dispersed when co-inoculated with motile cells. By teasing apart the abiotic and biotic dimensions, these 3D-printed devices will stimulate further research on microbial dispersal in soil and other water-unsaturated environments.

## Introduction

Spatial structure and physicochemical properties of microbial habitats strongly influence the composition and the ecological functions of microbial communities^1^. This is evident in water-unsaturated environments such as soils, where the combination of an unequal distribution of resources and the restricted movement of bacteria are often factors limiting their activity, as exemplified by the poor degradation of organic pollutants in soil ecosystems^2,3^. Limited dispersal is particularly significant in the case of bacteria, as most types of active dispersal require liquid films at the interface of biotic and abiotic surfaces^4,5^. Due to a lack of dispersal pathways in the form of continuous liquid films in water-unsaturated environments, the colonization and redistribution of bacteria to suitable habitats are restricted by their effective motility. In contrast to bacteria, dispersal limitation in water-unsaturated habitats is less of a problem for filamentous fungi^6^. Fungal mycelia extend readily in soils, colonizing also larger, air-filled pores that hinder bacterial dispersal most severely^7,8^. Fungal mycelia can reach up to a total length of 10^2^ m/g, 10^3^ m/g, and 10^4^ m/g in arable, pasture, and forest soils, respectively^9,10^, providing a highly efficient and ubiquitous three-dimensional connective scaffold.

In recent years, several studies have shown that motile bacteria can utilize the continuous liquid films on the surface of likely hydrophilic and semi-hydrophobic hyphal networks to disperse, using so-called “fungal highways” ^11–15^. Fungal networks are, however, not only a physical support resembling the highway system built by humans. Instead, hyphal networks continuously grow, reshape, and interact dynamically with the biotic and abiotic environment, including bacteria that disperse across them^16,17^. The diverse interactions between bacteria and fungi on these fungal highways remain difficult to study, partly due to the intertwining roles of fungi as both providers of a dispersal network and active biological players. For instance, one important aspect of fungal highways is that it requires bacteria to actively disperse on the network. This may occur using one of the motility types described so far: swimming and swarming, which rely on extracellular filamentous appendages, or other forms of dispersal such as crawling, gliding, or twitching^18^. However, to date, the relative importance of these different motility types to navigate on the fungal highway is still unknown. Also, several mechanisms occurring as a result of or triggered by fungal highways dispersal could be better studied if the physical and biological properties of the dispersal network could be teased apart. This includes investigating the role of chemotaxis^19^, cell-to-cell signaling^14,19,20^, germination of dormant cells^21^, gene transfer^22^, or co-metabolism^3,11,12,23^, all of which have been observed in previous studies involving fungal highways. While glass fiber networks have been previously used to study such interactions^8,12,24^, the rapid development of 3D-printing technologies, including miniaturization, provides us with the opportunity to make customized devices to study the physical component of fungal networks. By studying bacterial dispersal on 3D-printed models of fungal highways that provide a water film of defined properties (thickness, presence or absence of hydraulic flow), we can separate the part fungal mycelia play as a dispersal scaffold from the potential biotic interactions they exert on dispersing bacteria.

In this work, we designed two devices that reproduce a liquid-surface interface in the form of a meniscus, of varying thickness, as it may be found in hydrophilic soil surfaces or along fungal hyphae. The first device, which we term the “bacterial trail” can establish a relatively wide liquid film of 1.35 mm in lateral depth. The second device, which we term the “bacterial bridge” can establish much narrower liquid films of 0.14 mm in lateral depth. Motile bacterial cells dispersed in the liquid films in both devices. Non-motile cells were unable to disperse alone in the liquid film on the bacterial bridge device, but they could disperse (probably as “hitchhikers”^14,18,25,26^) in the presence of motile cells. Our devices lay a cornerstone for future investigations concerning microbial dispersal in natural unsaturated environments such as the soil, and can be used to investigate the diverse and dynamic interactions between soil microorganisms occurring on fungal highways^18,27^. They also serve as a proof of principle of the vast potential of applying 3D-printing technologies to rapidly design, prototype, and produce customized tools in modern biology labs and their particular application in microbial ecology^28^.

## Results

### Design and 3D-printing of the devices

We designed two devices in an attempt to represent fungal hyphae under different water-saturation conditions. Printed parts can nowadays reach a resolution of 50 μm on the *x-* and *y*-axes and a resolution between 25 μm to 100 μm on the *z*-axis^29^; this is however still far from the real dimensions of a fungal hypha (typically hyphae are reported to have diameters on the order of 1–30 μm^30^). To replicate the liquid film formed by surface tension next to a hypha, we relied on a bar-shaped structure printed with a hydrophilic resin capable of maintaining a continuous liquid meniscus when connected to liquid reservoirs. In the first device, the “bacterial trail” (hereafter “trail” for brevity), the bar is placed on top of eight discrete liquid reservoirs (Figure 1a); the “trail” device corresponds to wet conditions, where the liquid films forming on hydrophilic surfaces would be relatively thick (see below). The inoculation and end sampling wells were located directly below the bar along the longitudinal axis of the device. The other sampling wells (numbered from 1 to 6) were positioned to the sides of the bar, served to maintain a uniform thickness of the liquid film, and can also be used for sampling. In the liquid film on the “trail” device, small volume changes during inoculation or sampling can generate flow caused by pressure differences between the sampling wells, which may be able to transport non-motile particles such as bacterial cells and virus particles that do not have the ability to disperse actively. In the second device, the “bacterial bridge” (hereafter “bridge” for brevity), the bar mimicking fungal hyphae is placed at the top of two capillaries of 1 mm in diameter and 15 mm in height connected to two liquid reservoirs (interchangeable start and end sampling wells) (Figure 1b); this device corresponds to water-limiting conditions where the liquid films are thinner and less susceptible to disturbance by volumetric changes in the reservoir, despite being continuous along the hydrophilic surface^31,32^. In this way, the water film was also created and maintained by surface tension, but in contrast to the “trail” device, sampling or changes in the volume of the liquid in the reservoirs did not generate flow. The dimensions of different parts of the two devices are provided in the detailed computer-aided design (CAD) illustrations in Appendix Figure S1. The two devices were 3D-printed using a masked stereolithography (MSLA) printer. Corresponding Standard Tessellation Language -STL-files are provided in the ESM. The printing material has the desirable properties of being hydrophilic (with a contact angle of 52.8°, measured using a snake-based approach^33^), autoclavable, and biocompatible (Appendix Table S1 and Figure S2). Detailed printing and subsequent preparation procedures are provided in the Materials and Methods.

**Figure 1.**
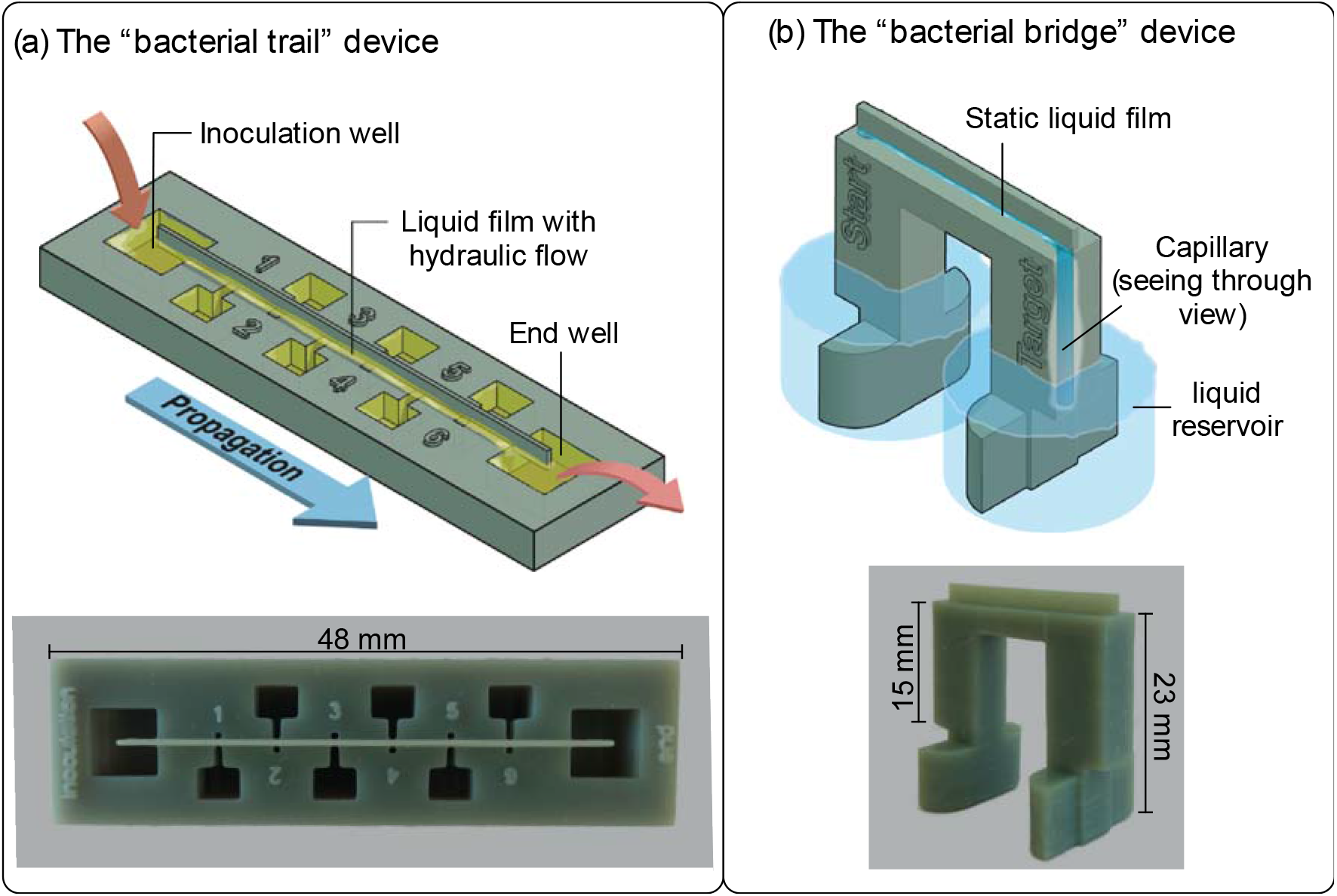
Cartoon illustrations and photos of the “bacterial trail” and “bacterial bridge” devices. The dimensions of different parts of the two devices are provided in the detailed CAD illustrations in Appendix Figure S1. In the “trail” device (panel a), a bar-shaped support structure connects eight liquid reservoirs, including an inoculation well, an end sampling well, and six intermediate sampling wells. In the “bridge” device (panel b), the support structure to form liquid films is located on the top of two capillaries of 1 mm in diameter and 15 mm in length.

### Characterization of the liquid films formed on the 3D-printed devices

We generated liquid films on the “trail” and “bridge” devices (Figure 2) following the procedures shown in ESM videos 1, 2, and described in the Materials and Methods. To control for the consistency of liquid film thickness (lateral depth), we measured the liquid films on four independently printed devices of each type at nine different locations on each device (Figure 2, Box-Whisker charts). The liquid film lateral depth did not differ significantly between devices of the same type by pairwise Z-tests with a threshold *p*-value of 0.0083 (six pairwise comparisons for each device) adjusted using the Bonferroni correction (ESM Table S2). Across the “trail” devices, the liquid film had a lateral depth of 1.35 ± 0.32 mm (*n*=4). Across “bridge” devices, the liquid film had a lateral depth of 0.14 ± 0.04 mm (*n*=4), approximately one-tenth of the liquid film lateral depth on the “trail” devices. The liquid film width on the “bridge” device can be adjusted by varying the length of liquid columns in the capillaries, which can be achieved by adding different volumes of liquid to the reservoirs (Appendix Figure S3). However, attention must be paid as the thinner the liquid film is, the more likely it will break during the duration of bacterial dispersal experiments if those last several days. The broader liquid film on the “trail” device allows flow caused by hydraulic pressure differences between the sampling wells. Using fluorescein as a marker, we measured the flow rate as the mean speed of marker transportation in the flow (1.19 mm/h, ESM Figure S2, Video 3). In contrast, the narrower liquid films on the “bridge” device and the vertical capillaries connecting the liquid films to the sampling wells effectively suppressed flow caused by disturbances to the liquid in the sampling wells. We found that the traveling speed of fluorescein from the start well to the end well can be explained by diffusion alone, suggesting the absence of hydraulic flow (ESM Figure S2).

**Figure 2.**
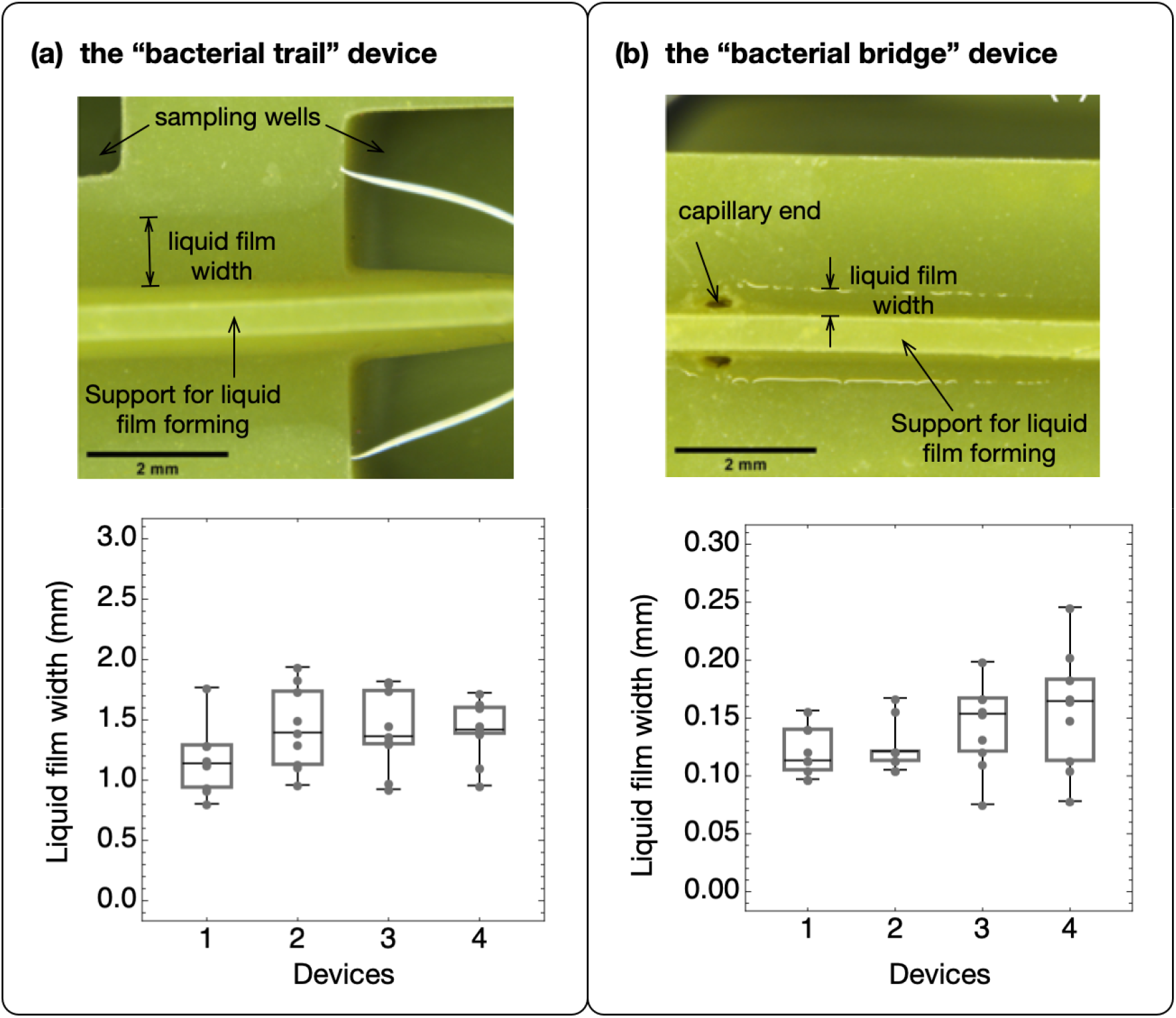
Stereoscopic pictures of the liquid film on the “bacterial trail” (panel a) and “bacterial bridge” (panel b) devices, and the distribution of the liquid film widths on four independently printed devices of each type. The photograph in each panel shows a typical stereoscopic image of the liquid films on the corresponding device. For each of the four devices of each type, a total of nine measurements of the liquid film width were measured and overlaid with the corresponding box-whisker chart. The middle line in each chart shows the median of the data points; the upper and lower boundaries of the boxes mark the 25% and 75% quantiles of the data. The liquid film widths do not differ significantly within a device type.

### Active and passive dispersal in the liquid films on the two 3D-printed devices

We used *Pseudomonas putida* KT2440 and its ⍰ *fliM* non-motile mutant strain^31^ to test the active and passive dispersal of cells in the liquid films on the “trail” and “bridge” devices.

#### Experiments with the “trail” device

Because of the presence of hydraulic flow in the liquid films on the “trail” device, we hypothesized that both strains could passively disperse from the inoculation well to other sampling wells driven by the flow. In addition, we expected the motile strains to have a dispersal advantage due to their ability to swim when sufficient energy sources are present in the environment. To test these hypotheses, we performed experiments under a full factorial design with the two bacterial strains (motile and non-motile), a culturing medium (nutrient broth (NB) that contains the required nutrients for active dispersal and cell replication, and phosphate-buffered saline (PBS) solution, which does not contain any usable energy source), and two dispersal durations (four hours and 24 hours), each with four replicates. At the start of the experiments, we inoculated 5 μl cell suspension (at an optical density of one) of either strain in the inoculation well. The cell density in each sampling well was quantified at the end of the corresponding dispersal duration (Figure 3a–b). To establish the relative contribution of growth versus dispersal, we compared the growth curves of the two strains in NB and PBS (Figure 3c).

**Figure 3.**
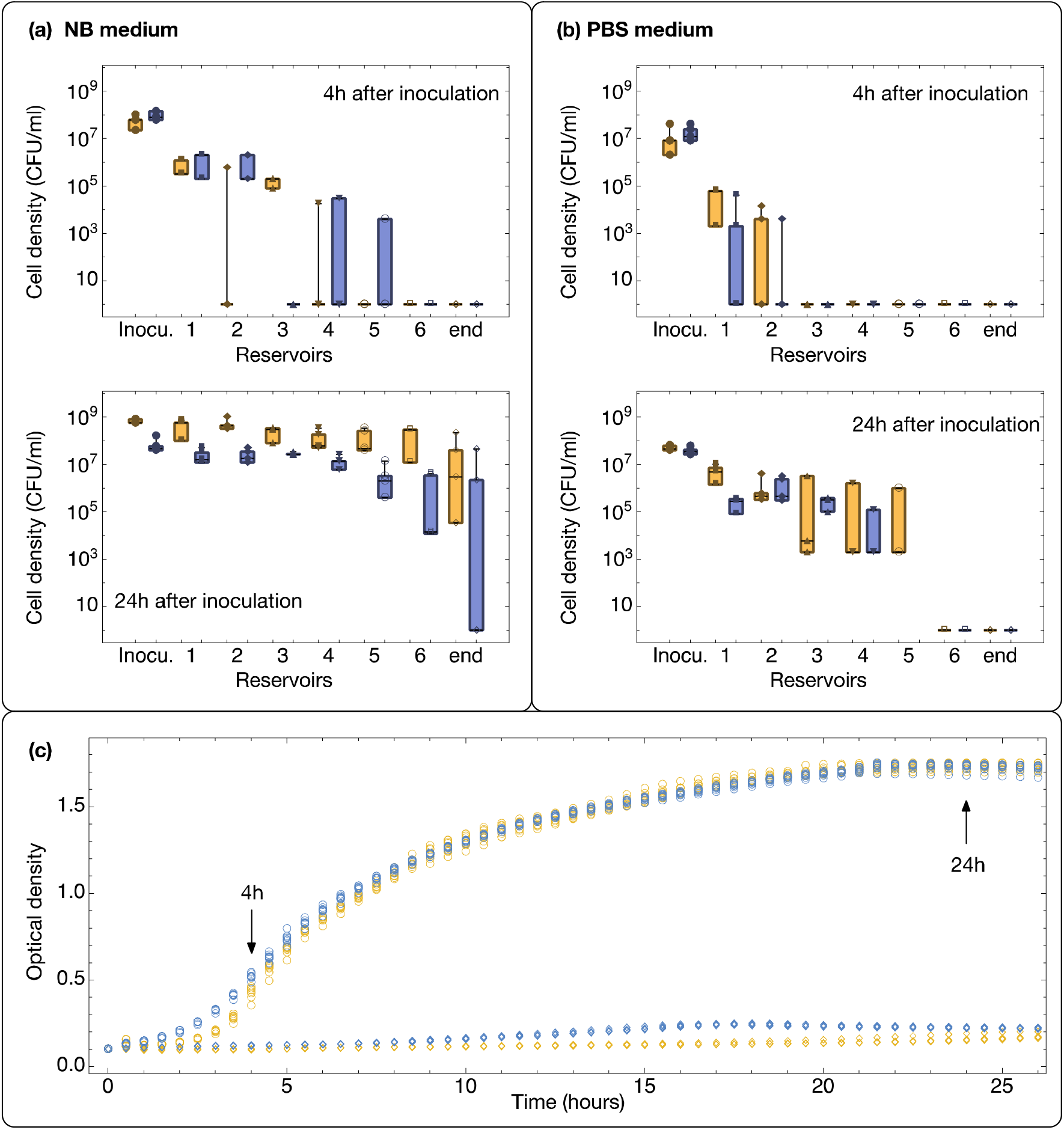
(a—b) Dispersal of motile (yellow) and non-motile (blue) bacterial strains on the bacterial “trail” device. Box-whisker charts show the density of bacterial cells in each of the sampling wells at the end of the corresponding dispersal duration: four hours (top panels) or 24 hours (bottom panels). Each box represents the data of four independent replicates. The data points are overlaid with the corresponding box-whisker chart. The devices were filled with two different liquid solutions: Nutrient Broth (NB) (panel a) and PBS buffer (panel b). For plotting on the log scale, we assigned a cell density of 1 CFU/ml to samples where no cells could be detected. (c) Growth curves of the motile (yellow) and non-motile (blue) strains in the NB medium (circles) and in PBS (diamonds). The plot is based on eight independent replicates for each bacterial strain and medium combination.

For both dispersal durations, there was a general trend of decreasing cell densities at larger distances to the inoculation well. Robust two-way ANOVA tests showed that the density of cells differed significantly across sampling wells after four hours (*p*=0.002 for in NB and *p=*0.06 in PBS) and after 24 hours (*p*<0.001 in NB and in PBS) of dispersal. The cell densities did not differ significantly between the motile and non-motile strains in either condition (*p=*0.299 in NB and *p=*0.585 in PBS) after four hours of dispersal. After 24 hours, the motile strain was significantly more abundant than the non-motile strain across sampling wells in NB (*p=*0.001) but not in PBS (*p=*0.1). In the NB medium, the growth curves of the motile and non-motile strains differed significantly (*p*<0.001) within the first four hours of growth, with the non-motile strain growing significantly faster than the motile strain, but the overall growth did not differ significantly (*p*=0.3) between the strains. In the PBS medium, both strains experienced minimal growth, with their optical densities barely doubled by 24 hours, although the non-motile strain grew consistently more than the motile strain (*p*<0.001).

#### Experiments with the “bridge” device

Because of the absence of hydraulic flow in the thin liquid film on the “bridge” device, we expected that only motile bacterial cells inoculated into the start well could disperse across the “bridge” and arrive at the target well. To test this hypothesis, we inoculated the motile and non-motile cells separately into the start well and tested the presence and density of cells in the target well after 72 hours. Our pilot experiments showed that the motile strain already arrived at the target well within 20 hours (data not shown), while the non-motile strain could not cross the “bridge” by diffusion in 72 hours. The culturing medium contained nutrients (PBS+1% NB+1% Ficoll) to allow for energy-consuming bacterial swimming, but sustained minimal growth and prevented the formation of a biofilm that could block the capillaries or help the non-motile strain to reach the target well through biofilm expansion (the growth curves of the two strains in the medium are provided in Appendix Figure S6). Our experiments showed that the motile cells arrived at the target well across all four replicates, while the non-motile cells did not arrive in all three replicates (Figure 4a).

**Figure 4.**
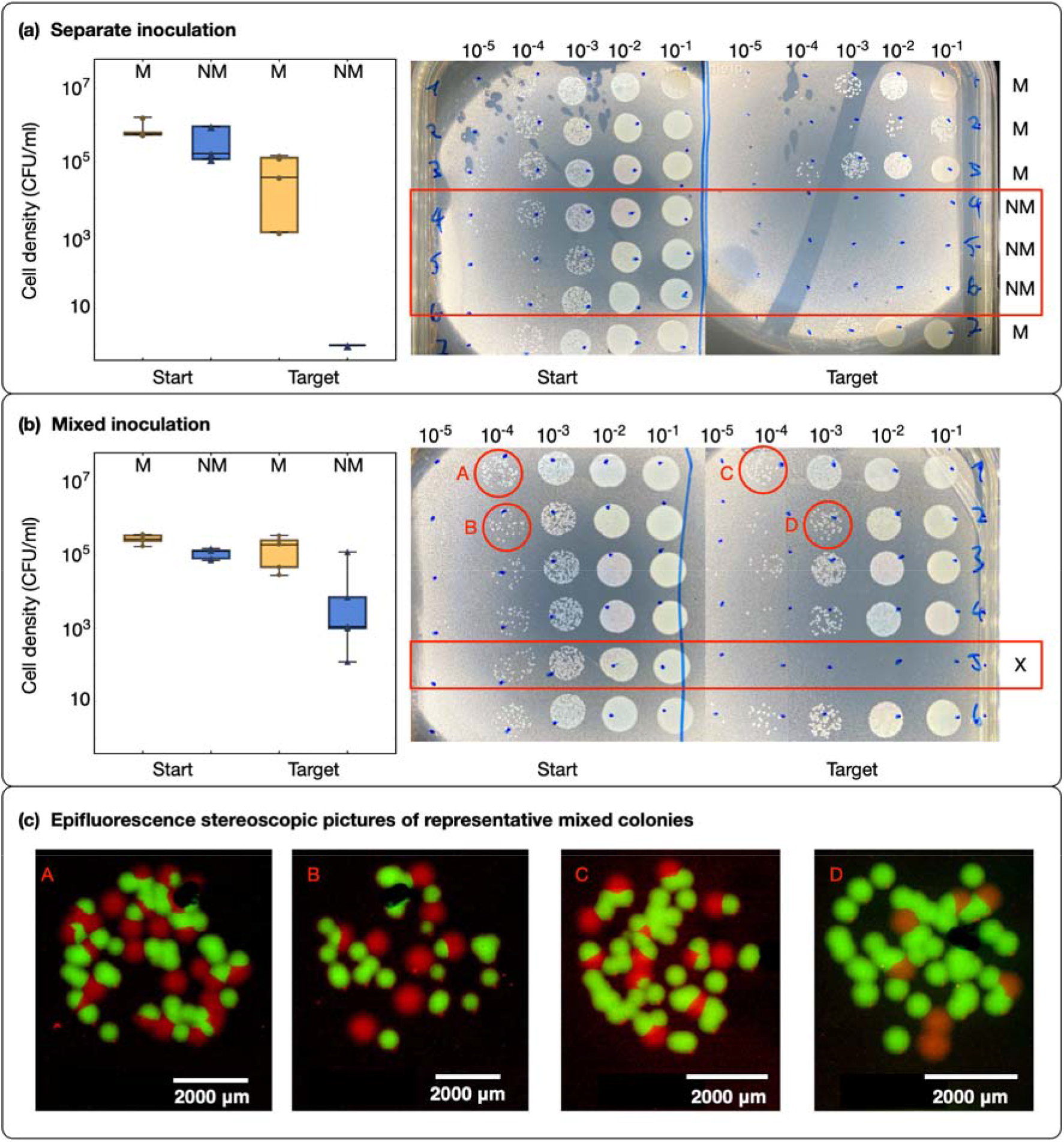
Active dispersal of cells on the “bacterial bridge” device. (a) The box-whisker chart shows the cell densities of the motile (M) and non-motile (NM) bacterial cells in the start well and the target wells at 96 h after inoculation. The two bacterial strains were inoculated separately with four replicates for the M strain and three replicates for the NM strain. The two photographs show the dilution series performed to quantify cell densities in the start and target wells for each replicate. The corresponding dilution ratio is shown on top of each column (increasing level of dilution from right to left). Replicates 1, 2, 3, and 7 correspond to the motile strain; replicates 4, 5, and 6 correspond to the non-motile strain (marked with a red box in the image). (b) The box-whisker chart shows the same information as in panel (a) except that the two bacterial strains were mixed at a 1:1 ratio immediately before inoculation. The two photographs show the dilution series for quantifying cell densities in the start and target wells. Replicate five (marked with X) did not produce reliable results because the liquid film on top of the bacterial bridge broke during the experiment. (c) Epifluorescence stereoscopic micrographs of the mixed colonies in circles A to D are presented in the corresponding figures in the lower role of the panel (b).

Non-motile bacteria are known to disperse along fungal hyphae by “hitchhiking” on motile bacterial cells^14^. To test whether this could happen in our system, we inoculated the motile and non-motile cells at a 1:1 ratio of cell numbers into the start well and tested the presence and cell densities of each cell type in the target well after 96 hours. Both motile and non-motile cells were detected in the target well in five out of six replicates (Figure 4b). In one replicate, the liquid film broke due to evaporation during the course of the experiments so that neither motile nor non-motile cells arrived at the target well (Figure 4b, the failed replicate 5 is marked with an X). Because the motile and non-motile cells are tagged with green and red fluorescent proteins, respectively, we could distinguish them upon plating (Figure 4c; corresponding color-blind-friendly pictures are provided in the ESM Figure S4).

## Discussion

### 3D-printed devices as models to investigate bacterial dispersal

The dispersal capacity of bacteria in water unsaturated environments plays a crucial role in determining the spatial and temporal dynamics of microbial communities and in turn their ecological functions^5^. In this work, we use 3D-printing to produce two devices that generate continuous liquid films in which bacteria can disperse. The variable thickness of the films produced mimic typical liquid films formed on hydrophilic biological (i.e., plant roots or fungal hyphae^12,14,15,34^) or abiotic surfaces. Thus, the devices can be used in control experiments involving bacterial dispersal on natural supports to help disentangle abiotic processes and biotic interactions. Due to the properties of the resin used, the devices have a hydrophilic surface for continuous liquid films to form, and are heat resistant up to 140 °C, enabling the devices to be autoclaved for sterilization and repeated use. The material cost for producing the devices is also low (0.35 USD per “bridge” device and 0.5 USD per “trail” device), allowing for affordability and widespread use.

The new experimental systems we developed allow for the study of bacterial dispersal under different water saturation conditions. Under some natural conditions, bacterial cells can be transported passively by flowing streams of water, such as percolation after rainfall or concentrated flow in cracks and macropores formed by growing plant roots in relatively dry soils ^35,36^ The relatively thick water films formed by the “trail” device serve to model this situation. By contrast, in unsaturated soils, bacterial dispersal is restricted by cell-substrate viscous drag and capillary pinning forces imposed by the thin and fragmented water films on the surface of substrates ^31,32^ In these cases, the pore water velocity drops below the movement speed of motile bacterial cells, and the effect of passive transportation (i.e., in hydraulic flow) on cell motion is then negligible^37^. The relatively thin and stationary water films formed by our “bridge” device capture this situation. A previous soil model showed that liquid films thicker than 5 μm were rare and rapidly drained even under mild suction conditions, and the resulting patchiness of the liquid films severely limits the dispersal capacity of bacteria^5^. Our results with the “bridge device” extended the conditions where non-motile bacteria cannot disperse alone in unsaturated soil. We showed that even in continuous water films (0.14 ± 0.04 mm on the “bridge” device) that were much broader than 5 μm, the non-motile cells were still unable to disperse unassisted (in the timeframe of the experiments, 72 hours), although motile cells could efficiently disperse (over 2.1 cm). Similar results were found using glass fibers to represent fungal hyphae^12^. Although an estimation of the liquid film thickness on the mycelia of soil fungi under conditions corresponding to typical unsaturated soils is currently inexistent, we expect it to be thinner than the liquid films established with the bridge device. If this is the case, the widespread fungal highways may provide a dispersal advantage exclusive to flagellated bacteria and the non-motile bacteria that can hitchhike the dispersal of their associated carriers. This would be in accordance with the findings of a dispersal advantage of motile bacteria in cheese rind^38^, and may contribute to explain the maintenance of costly flagella in soil bacteria^39^. By helping to tease apart the abiotic and biotic aspects of the interactions between bacteria and fungal mycelial networks, the devices we developed in this study open the door to many interesting future studies. For example, they can be used to identify and characterize fungi-secreted bioactive substances such as chemo-attractants, chemo-repellents, toxins, and signaling molecules.

### Impact of liquid film lateral depth in dispersal of motile and non-motile cells

In the relatively thick liquid films established in the “trail” device (1.35 ± 0.32 mm in lateral depth), both motile and non-motile cells can disperse passively, driven by hydraulic flow. This explains why there was no significant difference in dispersal between motile and non-motile cells in the PBS solution across sampling wells (Figure 3b). Because active dispersal by swimming with flagella requires energy^40,41^ and PBS lacks any carbon and energy sources, the motile cells can only disperse passively in a manner alike the non-motile cells. In the presence of energy sources, as in the NB medium, we expected the motile cells to have a dispersal advantage by being able to disperse actively by swimming, in addition to passive dispersal. Although the experimental results after four hours did not support our hypothesis with statistical significance (probably due to stochastic effects on the distribution of cells within a short duration of time and the higher growth rate of the non-motile cells in the first hours after inoculation, which counterbalanced the dispersal advantage of the motile strain), the experimental results after 24 hours provided clear evidence for a dispersal advantage of the motile cells over the non-motile cells, supporting our hypothesis.

In contrast to the relatively thick liquid films on the “trail” device, the liquid films on the “bridge” device are much thinner (0.14 ± 0.04 mm). The absence of hydraulic flow in these liquid films prevents non-motile cells from dispersal by drifting in the flow. This was shown in the single-strain inoculation experiments (Figure 3a), which confirmed the importance of the flagellum for dispersal on hydrated surfaces such as in the case of fungal highways^34,38,39^. Interestingly, non-flagellated bacteria were still able to disperse in the presence of actively swimming motile cells, as shown in the co-inoculation experiments (Figure 3b). Since the device does not introduce any biotic interaction between the dispersal network and the dispersing bacteria, we infer that the dispersal of non-motile bacteria facilitated by fungal highways^14,20^ may not require an active role of the fungus other than providing a continuous liquid pathway and a source of nutrients^21^. Although we do not know the exact mechanism explaining how the non-motile bacterial cells “hitchhike” to disperse in this particular case, existing literature provides some hypotheses that could be tested using our 3D-printed devices. For instance, an aspect that could be investigated in the future is whether physical attachment of the non-motile to the motile cells is required for transport or if the non-motile bacteria are dragged by the flow generated by motile bacteria. In previous studies, several non-motile bacterial species were found to attach to the cell body of the motile carrier bacterium *Capnocytophaga gingivalis*^42^; similarly non-motile bacterial and fungal spores were found to attach to the flagella of their corresponding carriers^15,25^. Based on these observations, different combinations of carriers and hitchhikers can be considered to test the possible release of chemicals to attract motile cells for attachment and dispersal^26^, as well as the transport of non-motile cells as useful “cargo” to degrade antibiotics^43^, or to invade an occupied niche^44^. These functional components of hitchhiking could be investigated in future experiments using our devices in combination with parallel experiments performed using fungal hyphae or other biological surfaces (e.g., plant roots).

### Limitations and opportunities of 3D-printed devices

Despite the advantages of the 3D-printed devices (i.e., fast prototyping, low cost, and reusability), there are still technological limitations that are relevant to mention. The most important limitation is the relatively low printing resolution, which leads to roughness with staircase-like patterns on the printed surfaces that are not parallel or perpendicular to the printing bed. This limitation prevented us from generating a highly obtuse angle between support structures with smooth surfaces to reduce the thickness of the liquid film on the devices, which may be achievable by other (often more expensive) microfabrication technologies such as photolithography, which has been used in producing microfluidic devices. In the case of the “bridge” device, the liquid columns in the vertical capillaries suppress hydraulic flow that can transport non-motile cells^37^. In principle, the thinner the capillaries, the better the effect of suppressing flow. However, the printing resolution set a lower limit on the diameter of the capillaries (0.5 to 1 mm in our case). Because the capillaries are perpendicular to the plane where the liquid films form (and we need to prioritize the smoothness of printing on that surface), it was not possible to further reduce the diameter of the capillaries (e.g., by reorienting the devices on the printing bed). The limit on capillary diameter may have contributed to the breakage of the liquid film in one of the six replicates in the bacterial hitchhiking experiment (Figure 4b). Despite this limitation, the bridge device is shown to be able to establish and maintain a thin water film of consistent width (Figure 2b) for experiments over long durations (96 hours in the hitchhiking dispersal experiments) with satisfying reliability. Moreover, the 3D-printed devices offer the possibility of investigating the effect of two different boundaries (i.e., a surface-water and a water-air boundary) that complements other approaches such as microfluidic devices in which the water-air boundary is often not considered. In the last years, the cost of 3D printers has been decreasing rapidly to a few hundred USD. The expanding applications in the medical (especially dental) industry have promoted the development of a spectrum of hydrophilic biocompatible resins with diverse properties. In light of the increasing accessibility to 3D printing and affordable material costs, we view the two devices we developed in this work as early examples of much broader future applications of 3D printing in modern biology labs. The accessibility to rapidly designed and produced customized devices at affordable prices will play an increasingly important role in democratizing science and advancing education and research with more rational use of limited resources, such as in areas with disturbed production and supply chains.

## Materials and Methods

### 3D printing and subsequent preparation of the devices

We designed the devices using the Autodesk Fusion 360 software, exported the design as STL files (provided in the ESM), prepared the files for printing with the PrusaSlicer v2.2.0 software, and printed them using the HTR-140 green resin (3DM) with a masked stereolithographic 3D printer (Prusa SL1). The devices were printed directly on the printing bed with a layer height of 0.05 mm at 17 °C. The exposure time for each layer was 40 seconds for the first layer and 15 seconds for the rest layers. After printing, the devices were washed, blown with compressed air, and then dried overnight at room temperature. Finally, the devices were UV cured in a Prusa CW1 curing and washing machine for 4 minutes.

### Building the liquid films on the devices and measuring their widths

To build a liquid film on the canal device, we first distributed 850 μl liquid (NB medium or PBS) in the eight sampling wells. Because of the hydrophilic property of the resin, water molecules can be attracted to the surface of the artificial hypha from the wells and spread along it automatically to form a liquid film. To make sure that the liquid film is continuous and is connected to each sampling well, we followed the artificial hyphae with the tip of a 1 ml pipette from one end to the other, going through all 8 wells. To build a liquid film on the bridge device, we first added 2.75 ml liquid each in two adjacent wells on a 24 well plate (Costar). We then placed the “start” and “target” ends of the bridge into the two wells. After that, we took 100 μl of liquid from the “start” well and ejected the liquid first at the openings of the two capillaries and then along the printed hypha. After a few seconds, most of the liquid flowed down along the two capillaries connecting the two sides of the “bridge” and the liquid surfaces in the two wells leveled off. The remaining liquid on top of the “bridge” device formed a continuous thin liquid film along the artificial hypha. Videos illustrating these procedures are provided in ESM videos 1 and 2.

The widths of the liquid film on each of the devices were measured by taking pictures with a stereoscope (Nikon SMZ18) with a scale bar as reference. We measured the liquid films on four independently printed devices of each type and took three pictures at different locations (one in the middle and two in proximity to either end of the artificial hypha) on each device. We then measured the width of the liquid film at three randomly selected positions in each picture. This resulted in nine independent measurements of the liquid film width for each piece of the devices. The pictures were analyzed using the Fiji software (version 1.8.0). Using the burned-in scale as a reference, we could measure the width of the liquid film. For each picture, we performed three measurements at different randomly selected positions, leading to a total of nine independent measurements of the liquid film width for each piece of the devices.

### Bacterial strains and the preparation for dispersal assays

We used an isogenic pair of motile and non-motile *P. putida* bacterial strains to test the active and passive transportation of cells in the liquid films formed on the devices. The motile KT2440 strain was isolated from the rhizosphere and is flagellated^45^. The non-motile strain is a non-flagellated (⍰ *fliM)* mutant of the KT2440 strain, obtained by allelic exchange with a truncated version of *fliM^31^.* The motile strain is tagged with the green fluorescent protein, and the non-motile strain is tagged with a red fluorescent protein (DsRed) marker. The two bacterial strains were cryo-preserved in 30% (v/v) glycerol at −80 °C. To reactivate the bacteria, cells were first plated on the Nutrient Agar (NA, 23 g/L, Carl Roth) through polygonal spreading and grown overnight at 30 °C in darkness. We then prepared bacterial suspensions from the overnight cultures by growing the cells in Nutrient Broth (NB, 23 g/L, Carl Roth) at ambient temperature under constant rotation at 120 rpm for 14 hours. The bacterial suspension was adjusted with NB as blank to an optical density of 1 (600 nm wavelength), and centrifugated at 5000 g for 5 minutes to collect cells. The collected cells were washed once and resuspended in 0.01 M phosphate-buffered saline (PBS: 1.5 g/L Na_2_HPO_4_·2H_2_O, 0.25 g/L NaH_2_PO_4_·2H_2_O, 8,5 g/L NaCl) with 1% NB as an energy source. The cell suspension of each strain was adjusted to a final OD of 1.

### Bacterial dispersal assays on the canal device

To test bacterial dispersal in the liquid film on the canal device, we performed experiments under a full factorial design with two different media (NB and PBS), two bacterial strains (motile and non-motile), and two dispersal durations (four hours and 24 hours), each with 4 replicates. After building the liquid films with either NB or PBS, we added 10 μl of the cell suspension of either strain to the inoculation well. To maintain the liquid film and prevent evaporation during the experiments, we placed the devices into a large glass Petri dish with four pieces of sterile filter paper with 1 ml of autoclaved deionized water added to each. We took a sample of 20 μl from each of the eight wells following the order from one to eight at the end of each assay. The cell density in each of the wells was quantified (see below).

### Bacterial dispersal assay on the bridge device

To test bacterial dispersal in the liquid film on the bridge device, we used a culture medium containing 0.01 M PBS with 1% of NB and 1% of Ficoll (10g/L, Sigma-Aldrich). We added Ficoll to the medium to increase its viscosity for suppressing flow^46^. As a thickening agent, Ficoll has the desirable properties of causing little osmotic stress and cannot be consumed or degraded by bacterial cells^47^. For the experiments, we first built liquid films on the devices and inoculated 5 μl of bacterial cell suspension (either the motile /non-motile strain alone or mixed with 1: 1 ratio) to the start well. The devices were kept in a closed Petri dish to keep humidity high in darkness for 72 hours for experiments where the inoculation contains only motile or non-motile cells. The experiments with inoculations that contain both motile and non-motile cells were run for 96 hours under the same condition. At the end of the experiments, we took a sample of 20 μl first from the target well and then from the start well of each device, and quantified the cell density in each sample.

### Cell density quantification

We performed a dilution series to quantify the cell density in each sample taken at the end of the dispersal assays. Each of the 20 μl samples were diluted five times by adding 180 μl NB (10x dilution by volume). We then inoculated 5 μl of each of the five dilutions on a rectangular NA plate with a multichannel pipette. The plates were sealed with parafilm and incubated at 30 °C in darkness for 24 hours. Afterwards, the lowest dilution that enabled counting of bacterial colonies formed by a single cell was chosen to determine the initial cell density in the corresponding sample.

## Supporting information

Appendix containing supplementary figures and tables

## Acknowledgement

We would like to thank Guillaume Cailleau for drawing the cartoon illustrations of the devices in Figure 1, and two anonymous reviewers for their constructive comments that helped us improve the paper. We also thank the Swiss National Science Foundation (grant PZ00P3_180145) and the U.S. Department of Energy (grant LANLF59T) for financial support.

