## Appendix containing supplementary figures and tables for "Design and construction of 3D-printed devices to investigate active and passive bacterial dispersal on hydrated surfaces"

#### Checklist of the ESM

1. Appendix with supplementary tables and figures
2. STL files of the printed devices
3. Supplementary video 1 — building liquid films on the "trail" device
4. Supplementary video 2 — building liquid films on the "bridge" device
5. Supplementary video 3 — transport of the fluorescein marker in the liquid films on the "trail" device
6. Supplementary video 4 — transport of the fluorescein marker in the liquid films on the "bridge" device

### Appendix

##### 1. Detailed design of the "bacterial trail" and "bacterial bridge" devices

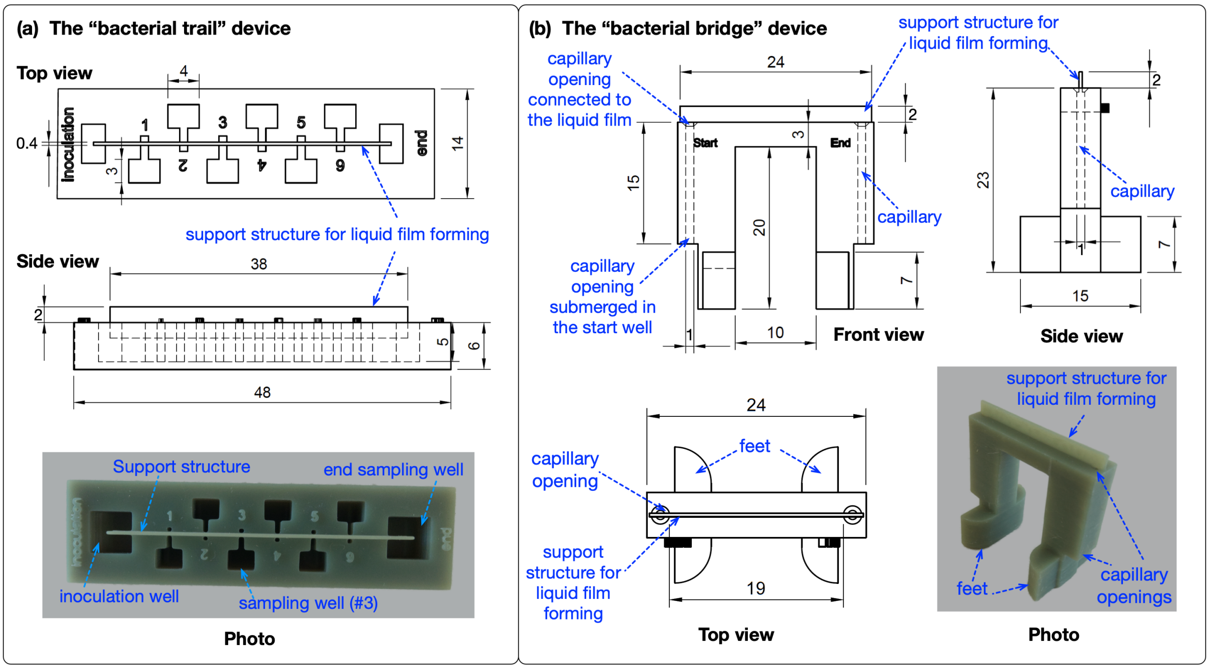


Figure S1. Design details and photographs of the "bacterial trail" and "bacterial bridge" devices. The detailed computer-aided design (CAD) illustrations indicate the dimensions of the components of the devices (corresponding Standard Tessellation Language -STL- files are provided in ESM). In the "trail" device (panel a), a bar-shaped support structure connects eight liquid reservoirs including an inoculation well, an end sampling well, and six intermediate sampling wells. In the "bridge" device (panel b), the support structure to form liquid films is located on the top of two capillaries of 1 mm in diameter and 15 mm in length. The lower openings of the capillaries next to the two feet of the device are submerged in liquid reservoirs in two wells on a 24-well plate. The upper openings of the capillaries connect to the liquid film formed along the artificial hypha on top of the device. All measurement values are in millimeters.

##### 2. Controlling for the biocompatibility of the printing material

To control for the biocompatibility of the printing material, we printed culturing tubes using the HTR-140 green resin (3DM) and cultured the bacteria *Pseudomonas putida* in the printed tubes and standard 50 ml

Falcon tubes (Sarstedt) of certified biocompatibility. We inoculated 10 µl of the bacterial suspension with an optical density equal to one into 5 ml NB medium and incubated the tubes in a thermos-shaker (Lab Gene Instruments) at 30 °C with a rotation speed of 120 r/min for 24 hours. There were three replicates for each treatment. Samples of 100 µl are taken at five time-points (immediately after inoculation, and 3, 6, 9, and 24 hours after inoculation) from each replicate. We measured the optical density of the samples to assess bacterial growth and compared the growth curves of bacteria in printed tubes and the standard Falcon tubes. Table S1 summarises the OD values at different time points. The growth curves in the printed tubes and the Falcon tubes are shown in Figure S1. We used the *compareTwoGrowthCurves* function of the *growthcurver* package in the R environment to compare the growth curves. The growth curves in printed and Falcon tubes did not differ significantly ($p$=0.54).

Table S1. Optical density of the culturing medium at five different time points after inoculating *P. putida* cells either in standard Falcon tubes or in printed tubes using the HTR-140 green resin.

| Treatment | Sampling time (hours after inoculation) | Mean optical density | SD of optical density |
|---|---|---|---|
| Falcon tube | 0 | 0.002 | 0 |
| Falcon tube | 3 | 0.010 | 0.0015 |
| Falcon tube | 6 | 0.0373 | 0.0050 |
| Falcon tube | 9 | 0.1053 | 0.0211 |
| Falcon tube | 24 | 0.5837 | 0.0811 |
| Printed tube | 0 | 0.002 | 0 |
| Printed tube | 3 | 0.011 | 0.001 |
| Printed tube | 6 | 0.0323 | 0.0076 |
| Printed tube | 9 | 0.083 | 0.0075 |
| Printed tube | 24 | 0.461 | 0.0650 |

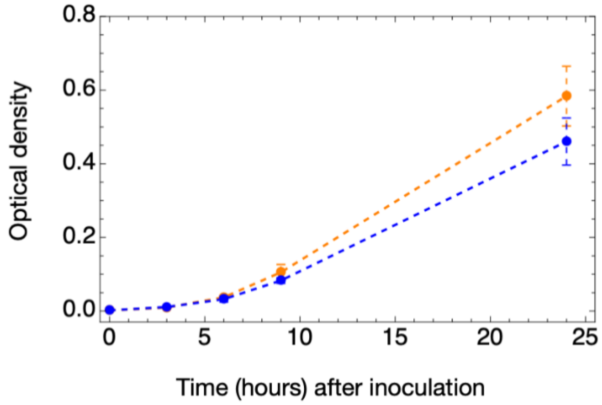


Figure S2. Growth curves of *P. putida* in biocompatible Falcon tubes (orange) and the printed tubes (blue).

**3. Adjusting the liquid film width on the "bacterial bridge" device**

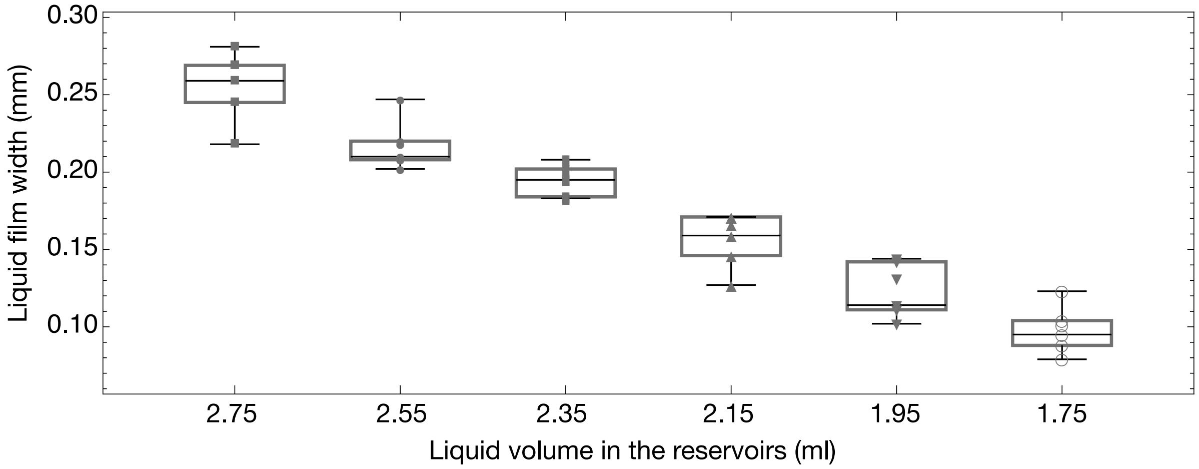


Figure S3. The liquid film width on the "bacterial bridge" device can be adjusted by varying the length of liquid columns inside the capillaries, which can be easily achieved by adjusting the volume of liquid added to the reservoirs. The results were produced using the wells on a standard 24-well plate as liquid reservoirs. However, attention must be paid as the thinner the liquid film is, the more likely it will break during the duration of bacterial dispersal experiments if those last several days.

**4. Controlling for the consistency of liquid film width across devices of the same type.**

To test whether different pieces of the same type of device produce liquid films of comparable width (Box-Whisker charts of the distributions are provided in Figure 2 of the main text), we performed pairwise Z-tests of the liquid film width distributions. To account for multiple comparisons (6 different pairwise comparisons for each device type), we use the Bonferroni correction to adjust the threshold *p*-value 0.05 to 0.00833. All the *p*-values of the pairwise comparisons are higher than the adjusted threshold (Table S2), and therefore, we conclude that the distributions of liquid film width are consistent across different pieces of devices of the same type.

Table S2. *P*-values of pairwise *Z*-tests of the liquid film width on four independently printed pieces of devices of each type.

| Device Type | Device pairs | *p*-value of pairwise Z-test |
|---|---|---|
| **Canal** | #1 vs #2 | 0.066 |
| | #1 vs #3 | 0.081 |
| | #1 vs #4 | 0.047 |
| | #2 vs #3 | 0.901 |
| | #2 vs #4 | 0.859 |
| | #3 vs #4 | 0.969 |
| **Bridge** | #1 vs #2 | 0.657 |
| | #1 vs #3 | 0.118 |
| | #1 vs #4 | 0.054 |
| | #2 vs #3 | 0.219 |

| | | |
|---|---|---|
| | #2 vs #4 | 0.094 |
| | #3 vs #4 | 0.511 |

**5. Transport of the fluorescein marker in the liquid films on the devices**

To determine whether hydraulic flows were present in the liquid films on the “bacterial trail” and “bacterial bridge” devices, we measured the transport speed of fluorescein in the liquid films. To do so, we added 10 µl fluorescein into the inoculation well on one “bacterial trail” device (Figure S4a) and three replicates of the “bacterial bridge” devices (Figure S4b). The transport speed of fluorescein on the “bacterial trail” device was faster than that on the “bacterial bridge” devices. The temporal dynamics of fluorescein transport on the “bacterial bridge” devices fit well to a one-dimensional diffusion model (Figure S4c, gray line), suggesting the absence of hydraulic flow in the liquid film.

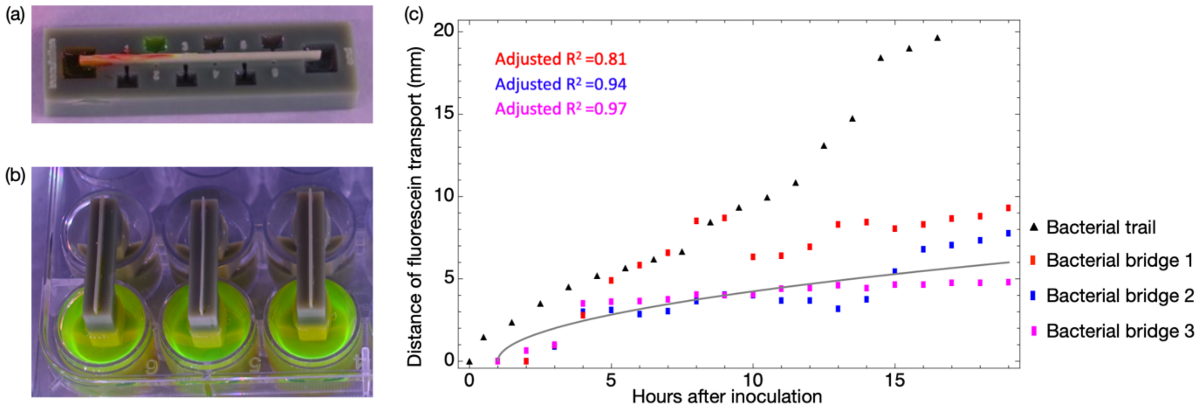


Figure S4. Transport of the fluorescein in the liquid film on the “bacterial trail” and “bacterial bridge” devices. (a) Photo of the “bacterial trail” device 10 hours after inoculating 10 µl of fluorescein into the inoculation well. (b) Photo of three replicates of the “bacterial bridge” devices immediately after inoculating 10 µl of fluorescein into the start wells. (c) Transport distance of the fluorescein over time in the liquid films on the devices. The gray solid line represents the expected diffusion speed of the fluorescein in the absence of hydraulic flow fitted from the one-dimensional diffusion model. The fluorescein did not start to diffuse in the liquid films on the “bacterial bridge” devices immediately after inoculation because we inoculated the fluorescein into the start well, and it took around one hour for the fluorescein molecules to reach the top of the capillaries that connect the liquid films to the corresponding start well.

**6. Adjusted main text Figure 4 for readers who could not distinguish red and green colors**

The motile and non-motile bacterial strains we used in the experiments were labeled with green and red fluorescent proteins, respectively. To facilitate the accessibility for readers with limited color vision, we provide below modified Figure 4c, where the red color was replaced by magenta. We further tested the result of color replacement with Fiji’s colorblindness simulator (version 1.8.0).

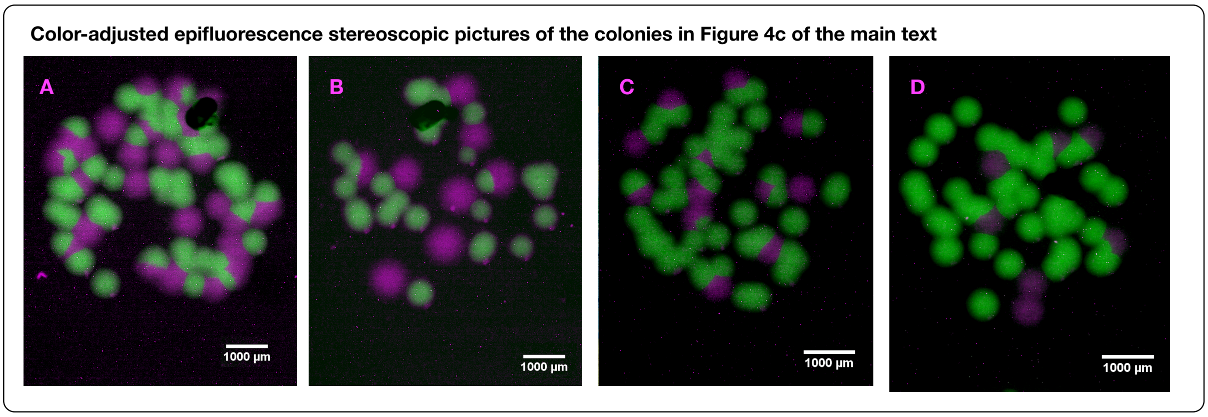


Figure S5. Color-adjusted Figure 4 panel c of the main text. The red color is replaced by magenta to distinguish it from the green fluorescence signal.

#### 7. Growth curves of the bacterial strains in different culturing media

To disentangle the contributions of growth and dispersal in the dispersal assays using the "trail" and "bridge" devices, we performed growth curves of the two strains in the three media we used in the experiments. In the experiments with the "trail" device, we used PBS and NB media. In the experiments with the "bridge" device, we PBS + 1% NB+ 1% Ficoll.

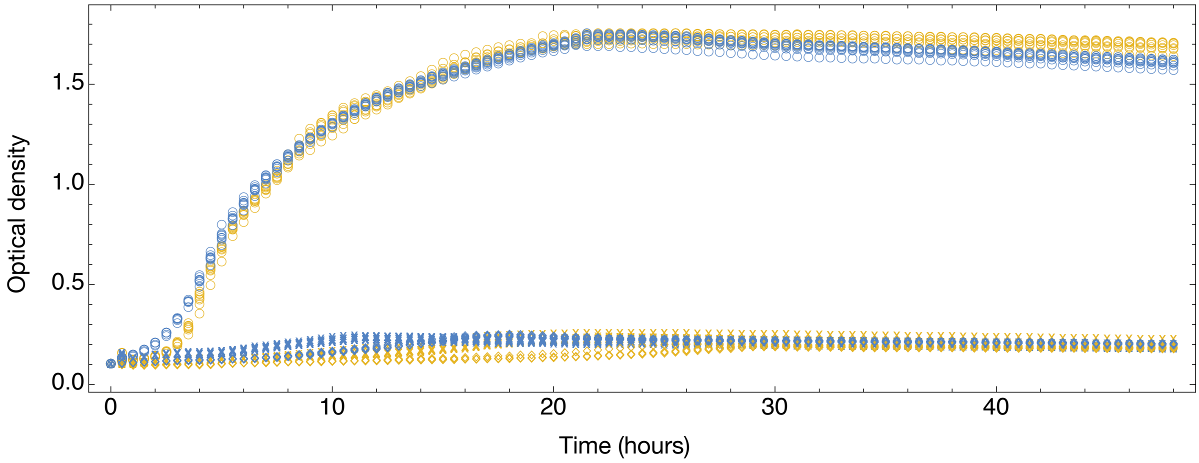


Figure S6. Growth curves of the motile (yellow) and non-motile (blue) strains in the NB medium (circles), PBS buffer (diamonds), and PBS + 1% NB + 1% Ficoll medium (crosses). The plot is based on eight independent replicates for each bacterial strain and medium combination.
